# Rapid Inactivation of SARS-CoV-2 by Silicon Nitride, Copper, and Aluminum Nitride

**DOI:** 10.1101/2020.06.19.159970

**Authors:** Giuseppe Pezzotti, Eriko Ohgitani, Masaharu Shin-Ya, Tetsuya Adachi, Elia Marin, Francesco Boschetto, Wenliang Zhu, Osam Mazda

## Abstract

**Introduction:** Viral disease spread by contaminated commonly touched surfaces is a global concern. Silicon nitride, an industrial ceramic that is also used as an implant in spine surgery, has known antibacterial activity. The mechanism of antibacterial action relates to the hydrolytic release of surface disinfectants. It is hypothesized that silicon nitride can also inactivate the coronavirus SARS-CoV-2.

**Methods:** SARS-CoV-2 virions were exposed to 15 wt.% aqueous suspensions of silicon nitride, aluminum nitride, and copper particles. The virus was titrated by the TCD_50_ method using VeroE6/TMPRSS2 cells, while viral RNA was evaluated by real-time RT-PCR. Immunostaining and Raman spectroscopy were used as additional probes to investigate the cellular responses to virions exposed to the respective materials.

**Results:** All three tested materials showed >99% viral inactivation at one and ten minutes of exposure. Degradation of viral RNA was also observed with all materials. Immunofluorescence testing showed that silicon nitride-treated virus failed to infect VeroE6/TMPRSS2 cells without damaging them. In contrast, the copper-treated virus suspension severely damaged the cells due to copper ion toxicity. Raman spectroscopy indicated differential biochemical cellular changes due to infection and metal toxicity for two of the three materials tested.

**Conclusions:** Silicon nitride successfully inactivated the SARS-CoV-2 in this study. The mechanism of action was the hydrolysis-mediated surface release of nitrogen-containing disinfectants. Both aluminum nitride and copper were also effective in the inactivation of the virus. However, while the former compound affected the cells, the latter compound had a cytopathic effect. Further studies are needed to validate these findings and investigate whether silicon nitride can be incorporated into personal protective equipment and commonly touched surfaces, as a strategy to discourage viral persistence and disease spread.

## 1. Introduction

The novel coronavirus, SARS-CoV-2, has led to a worldwide pandemic and raised interest in the surface-mediated transmission of viral diseases [1–3]. Respiratory aerosols and droplets, and contaminated surfaces facilitate viral spread from person to person leading to recommendations of social distancing, wearing of masks, hand-washing, and regular surface disinfection. Data suggest that the SARS-CoV-2 virus can remain viable on copper, plastic, steel, and cardboard surfaces for 4-72 hours after contact, and up to 7-days on surgical masks [4]. Viral persistence on these and other materials presents a risk for the social and nosocomial propagation of COVID-19, the disease caused by SARS-CoV-2. Current viral inactivation methods include the surface application of chemicals, such as a combination of ethanol with hydrogen peroxide or sodium hypochlorite [2]. Irradiation of surfaces with ultraviolet light is another virus disinfection strategy [5]. These and other proposed antiviral methodologies are ultimately limited by their toxicity to human cells [6,7]. As a practical solution, surfaces that are safe for human contact and capable of spontaneously inactivating viruses are desirable to control the spread of viral diseases.

Silicon nitride (Si_3_N_4_) is a non-oxide ceramic that has been used in many industries since the 1950s. A formulation of Si_3_N_4_ is FDA-cleared for use as an intervertebral spinal spacer in cervical and lumbar spine fusion surgery, with proven long-term safety, efficacy, and biocompatibility. Clinical data for Si_3_N_4_ implants compare favorably with other spine biomaterials, such as allograft, titanium, and polyetheretherketone [8–13]. A curious finding is that Si_3_N_4_ implants have a lower incidence of bacterial infection (*i*.*e*., less than 0.006%) when compared to other implant materials (2.7% to 18%) [14]. This property reflects the complex surface biochemistry of Si_3_N_4_ that elutes minute amounts of nitrogen, which is converted to ammonia, ammonium, and other reactive nitrogen species (RNS) that inhibit bacteria [15]. A recent investigation also found that viral exposure to sintered Si_3_N_4_ powders in aqueous suspension inactivated H1N1 (Influenza A/Puerto Rico/8/1934), Feline calicivirus, and Enterovirus (EV-A71) [16]. Based on these findings, it is hypothesized that Si_3_N_4_ may be able to inactivate SARS-CoV-2.

The present study compared the effects of exposing SARS-CoV-2 to aqueous suspensions of Si_3_N_4_ and aluminum nitride (AlN) particles and two controls, (*i*.*e*., a suspension of copper (Cu) particles (positive control) and a sham treatment (negative control)). Cu was chosen as a positive control because of its well-known ability to inactivate a variety of microbes, including viruses [17]. AlN was included in the testing because, like Si_3_N_4_, it is a nitrogen-based compound whose surface hydrolysis in aqueous solution leads to the elution of ammonia, with an attendant increase in pH. Since comparable antiviral and antibacterial phenomena are believed to be operative for all nitride-based compounds, AlN was used to provide additional insight into the antipathogenic mechanisms of nitrogen-containing inorganic materials [18,19].

## 2. Materials and Methods

### Preparation of Test Materials

Si_3_N_4_, Cu, and AlN powders were acquired from commercial sources (SINTX Technologies, Inc., Salt Lake City, UT USA, FUJIFILM Wako Pure Chemical Corporation, Osaka, Japan, and Tokuyama Co., Yamaguchi, Japan, respectively). Si_3_N_4_ powder (nominal composition of 90 wt.% Si_3_N_4_, 6 wt.% Y2O3, and 4 wt.% Al2O3) [20] was prepared by aqueous mixing and spray-drying of the inorganic constituents, followed by sintering of the spray-dried granules (∼1700°C for ∼3 h), hot-isostatic pressing (∼1600°C, 2 h, 140 MPa in N_2_), aqueous-based comminution, and freeze-drying. The resulting powder had an average particle size of 0.8 ± 1.0 μm. As-received Cu powder (USP grade 99.5% purity) granules were comminuted to achieve a particle size comparable to the Si_3_N_4_. AlN powder had an average particle size of 1.2 ± 0.6 μm as received, which was comparable to Si_3_N_4_.

### Cells and Virus

VeroE6/TMPRSS2 cells (Japanese Collection of Research Biosources Cell Bank, National Institute of Biomedical Innovation, Osaka, Japan) were used in the viral assays [21]. Cells were grown in Dulbecco’s modified Eagle’s minimum essential medium (DMEM) (Nissui Pharmaceutical Co. Ltd., Tokyo, Japan) supplemented with G418 disulfate (1 mg/ml), penicillin (100 units/mL), streptomycin (100 μg/mL), 5% fetal bovine serum, and maintained at 37°C in a 5% CO2 / 95% in a humidified atmosphere. The SARS-CoV-2 (Japan/AI/I004/2020; Japan National Institute of Infectious Diseases, Tokyo, Japan) viral stock was propagated using VeroE6/TMPRSS2 cells at 37°C for 2 days.

### Viral infection

Fifteen weight percent (15 wt.%) of the Si_3_N_4_, Cu, and AlN powders were separately dispersed in 1 mL of PBS(-), followed by the addition of the viral suspension (2 × 10^5^ median tissue culture infectious dose (TCID_50_) in 20 μL). Due to the higher density of the Cu powder, its volumetric fraction was approximately one-third of the Si_3_N_4_. Mixing was gently performed at room temperature for 1 min by slow manual rotation or for 10 min using a rotation machine. After exposure, the powders were pelleted by centrifugation (2400 RPM 2 mins) followed by filtration through a 0.22 μm filter (Hawach Sterile PES Syringe Filter, HAWACH SCIENTIFIC CO., LTD., Xi’an, China). Supernatants were collected, aliquoted, and subjected to TCID_50_ assays and real-time RTPCR.

### Titration of virus

Experiments were performed in triplicate including sham-treated virus suspension that was not exposed to any powder. A confluent monolayer of VeroE6/TMPRSS2 cells in a 96-well plate was inoculated with 50 μL/well of each virus suspension in a tenfold serial dilution with 0.5% FBS DMEM (*i*.*e*., maintenance medium). Viral adsorption at 37°C for 1 h was made with gently shaking every 10 min. Afterward, 50 μL/well of the maintenance medium was added. The plate was incubated at 37°C in a 5% CO2 /95% humidified atmosphere for 4 days. The cytopathic effect (CPE) of the infected cells was observed under a phase-contrast microscope. The cells were subsequently fixed by adding 10 μL/well of glutaraldehyde followed by staining with 0.5% crystal violet. The TCID_50_ was calculated according to the Reed-Muench method [22].

### Viral RNA Assay

After exposure to the powders, 140 μL of the supernatants were used for viral RNA extraction. RNA was also extracted from the surfaces of the centrifuged and filtered powders. RNA purification was performed by using a QIAamp Viral RNA Mini kit (QIAGEN, Germantown, MD, USA). An aliquot of 16 μL of isolated RNA was reverse-transcribed using ReverTra Ace^®^ qPCR RT Master Mix (Toyobo, Shiga, Japan). Quantitative real-time PCR was performed using a Step-One Plus Real-Time PCR system (Applied Biosystems, Foster City, CA, USA) and two sets of primers/probes specific for viral N gene.

Set 1: (forward primer, 5’-CACATTGGCACCCGCAATC-3’; reverse primer, 5’-GAGGAACGAGAAGAGGCTTG-3’’;probe, 5’-(FAM) ACTTCCTCAAGGAACAACATTGCCA (BHQ) - 3’); and Set 2: (forward primer, 5’-AAATTTTGGGGACCAGGAAC-3’; reverse primer, 5’-TGGCAGCTGTGTAGGTCAAC-3’’; probe, 5’-(FAM) ATGTCGCGCATTGGCATGGA (BHQ) - 3’).

Each 20 μL reaction mixture contained 4 μL of cDNA, 8.8 pmol of each primer, 2.4 pmol of the probe, and 10 μL GoTaq Probe qPCR master mix (Promega, Madison, WI, USA). The amplification protocol consisted of 50 cycles of denaturation at 95°C for 3 s and annealing and extension at 60°C for 20 s.

### Immunofluorescence

Vero E6/TMPRSS2 cells on cover glass were inoculated with 200 μL of virus supernatant. After viral adsorption at 37°C for 1 h, the cells were incubated with the maintenance medium in a CO_2_ incubator for 7 h. For the detection of infected cells, they were washed with TBS (20 mM TrisHCl pH 7.5, 150 mM NaCl) and fixed with 4% PFA for 10 min at room temperature (RT) followed by membrane permeabilization with 0.1% Triton X in TBS for 5 min at RT. The cells were blocked with 2% skim milk in TBS for 60 min at RT and stained with anti-SARS Coronavirus envelope (Rabbit) antibody (Dilution =1:100) (ProSci Inc., Poway, CA, USA) for 60 min at RT. After washing with a buffer, cells were incubated with an Alexa 594 Goat Anti-Rabbit IgG (H+L) (1:500) (Thermo Fisher Scientific, Waltham, MA, USA) and Alexa 488 Phalloidin (1:50) (Thermo Fisher Scientific, Waltham, MA, USA) for 60 min at RT in the dark. ProLongTM Diamond Antifade Mountant with DAPI (Thermo Fisher Scientific) was used as a mounting medium. The staining was observed under a fluorescent microscope BZX710 (Keyence, Osaka, Japan). The total cells, Phalloidin-staining cells, and protein-expressing cells were counted using the Keyence BZ-X Analyzer.

### Raman Spectroscopy Assay

Vero E6/TMPRSS2 cells were infected with 200 μL of each virus suspension on glass sites. After viral adsorption at 37°C for 1 h, the infected cells were incubated with the maintenance medium in a CO_2_ incubator for 4 h and fixed with 4% paraformaldehyde for 10 min at RT. After washing with distilled water twice, infected cells were air-dried and *in situ* analyzed using a highly sensitive Raman spectroscope (LabRAM HR800, Horiba/Jobin-Yvon, Kyoto, Japan) with a 20× optical lens. It operated in microscopic measurement mode with confocal imaging in two dimensions. A holographic notch filter within the optical circuit was used to efficiently achieve a spectral resolution of 1.5 cm^-1^via a 532 nm excitation source operating at 10 mW. Raman emissions were monitored using a single monochromator connected to an air-cooled charge-coupled device (CCD) detector (Andor DV420-OE322; 1024 × 256 pixels). The acquisition time was fixed at 10 s. Thirty spectra were collected and averaged for each analysis time-point. Raman spectra were deconvoluted into Gaussian–Lorentzian sub-bands using commercially available software (LabSpec 4.02, Horiba/Jobin-Yvon, Kyoto, Japan).

### Statistical Analysis

The Student’s *t*-test determined statistical significance for n=3 and at a *p*-value of 0.01 using Prism software (GraphPad, San Diego, CA USA).

## 3. Results

### Median Tissue Culture Infection Dose

TCID_50_ assay results for the 15 wt.% Si_3_N_4_, Cu, and AlN powders are shown in Figure 1. Inactivation times of 1 and 10 mins are shown in Fig. 1(a)∼(b) and Fig. 1(c)∼(d), respectively. Relative to the negative control, all three powders were effective in inactivating SARS-CoV-2 virions (>99%) for the two exposure times.

**Figure 1.**
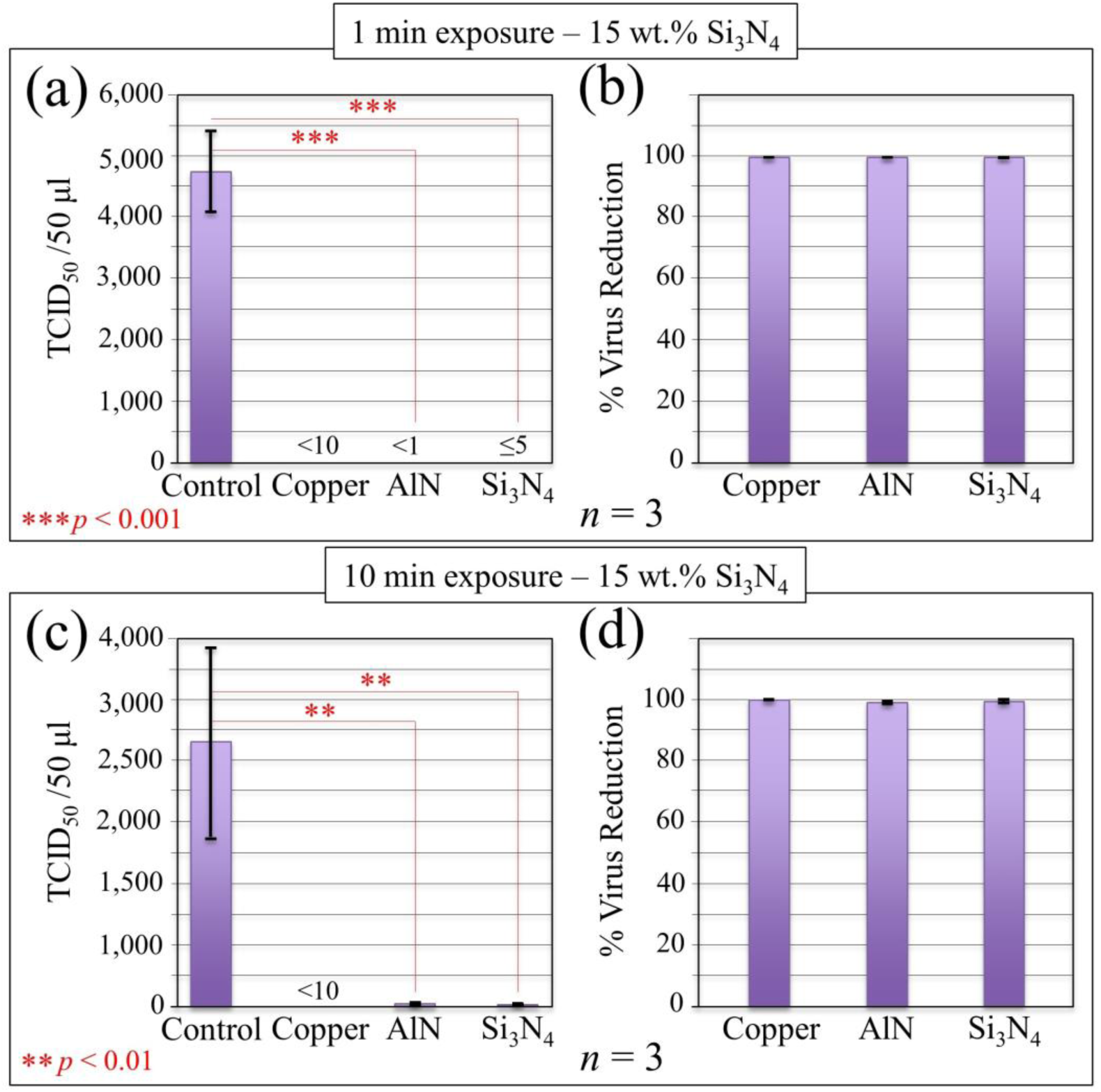
Inactivation of SARS-CoV-2 by nitride powders; virus suspensions were treated with 15 wt.% Cu, AlN, and Si_3_N_4_ powders in an aqueous medium at room temperature for 1- and 10-min. The control virus was treated identically without the addition of any powder. After light centrifugation and filtration, supernatants were subjected to TCID_50_ assay. The Reed-Muench method was used to determine the virus titers. TCID_50_/50 μL (a)/(b) and % reduction (c)/(d) are shown for virus inactivation times of 1 and 10 min, respectively. Statistics are given in the inset according to unpaired two-tailed Student’s t-test (n=3).

### RNA Fragmentation

To examine whether viral RNA was fragmented from exposure to the powders, RT-PCR tests were conducted on the viral N gene sequence. The results are shown in Fig. 2(a)∼(b) and (c)∼(d) for 1- and 10-min exposures, respectively. Again, in comparison to the supernatant of the negative control that was not exposed to any powder, almost complete fragmentation of the RNA was observed for Cu while significant damage was caused by AlN and to a lesser extent by Si_3_N_4_. After 10 min exposure to the powders, substantial cleavage of the RNA was seen for all three materials. While Cu still showed the most fragmentation, Si_3_N_4_ demonstrated similar effectiveness, and AlN was essentially identical to the 1-min exposure condition.

**Figure 2.**
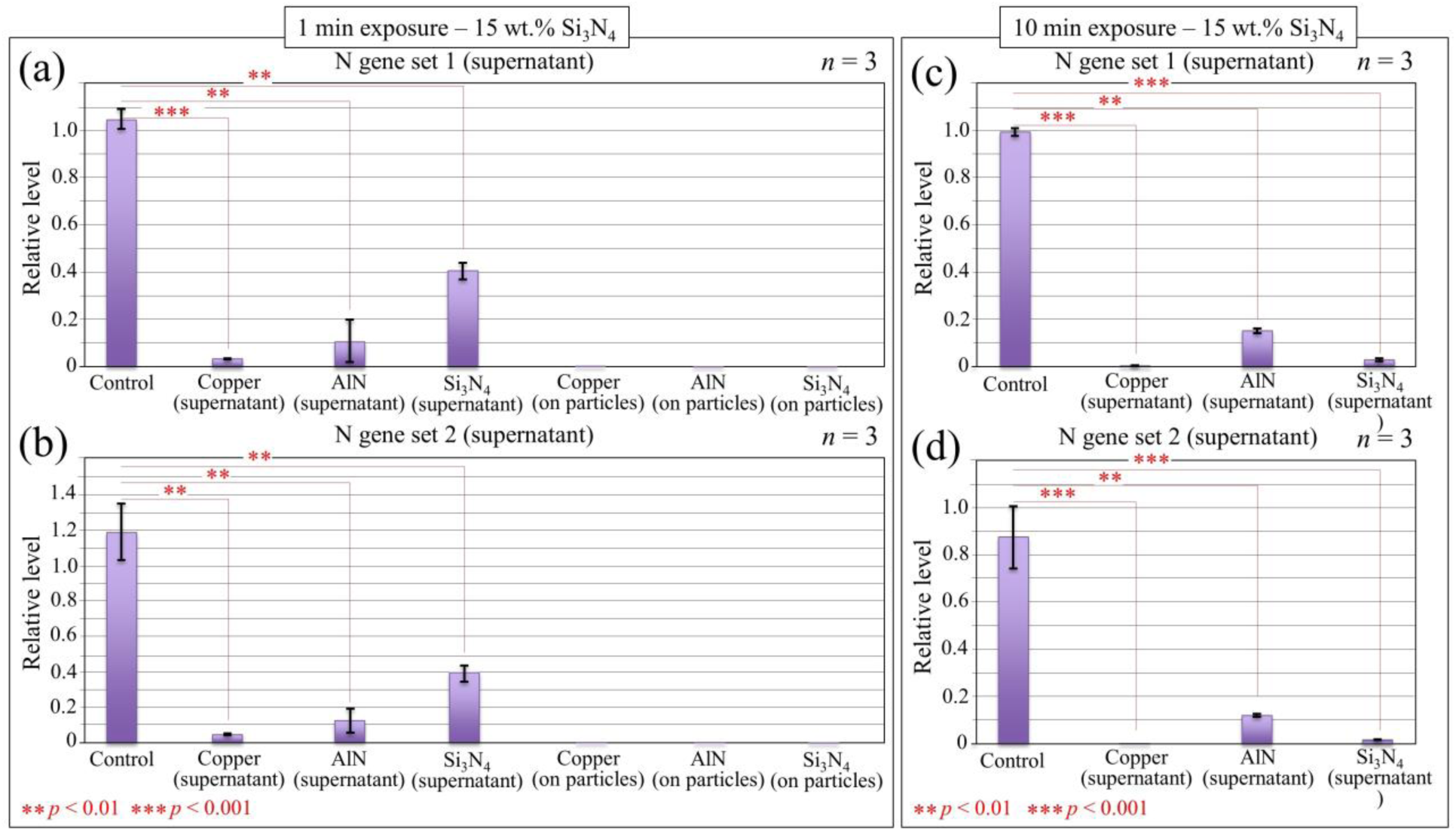
Viral RNA underwent severe degradation after exposure to copper or nitride particles. In (a) and (b), virus suspensions were exposed to Cu, AlN, and Si_3_N_4_ powders for 1 min, and viral RNA in supernatants and on particles were evaluated using viral N gene “set 1” and “set 2” primers, respectively. Data collected on supernatants and pellet samples are given in comparison with the amount of viral N gene RNA in suspension that was left untreated. In (c) and (d), results of RT-PCR tests for supernatants after 10 min exposure of virus suspension to Cu, AlN, and Si_3_N_4_ powders for viral N gene “set 1” and “set 2” primers are shown, respectively. Statistics are given in the inset according to unpaired two-tailed Student’s *t*-test (*n*=3).

Viral RNA was virtually undetectable for all three materials from the pelleted powders after 1 min of exposure (*cf*. Fig. 2(a)∼(b)). This result suggests that the decrease of viral RNA in the supernatant was not because of adherence of the RNA to the powders, but rather due to degradation.

### Immunofluorescence Testing

Immunofluorescence imaging using anti-SARS coronavirus envelope antibody (red), Phalloidin that stains F-actin in viable cells (green), and DAPI for cell nuclear staining (blue) was then used to confirm the TCID_50_ assay and gene fragmentation results. Figures 3(a)∼(d) show fluorescence micrographs representative of the VeroE6/TMPRSS2 cell populations that were inoculated with supernatants of (a) unexposed virions (*i*.*e*., negative control) and 10-min-exposed virions of (b) Si_3_N_4_, (c) AlN, and (d) Cu. Figure 3(e) shows cells that were not inoculated with the virus (labeled as “sham-infected” cells. The red-fluorescent signals in the negative control (Fig. 3(a)) demonstrated that the virus had infected the cells and viral protein was synthesized. This contrasts with the sham-infected cells (Fig. 3(e)), which did not show any evidence of expression of the viral protein.

**Figure 3.**
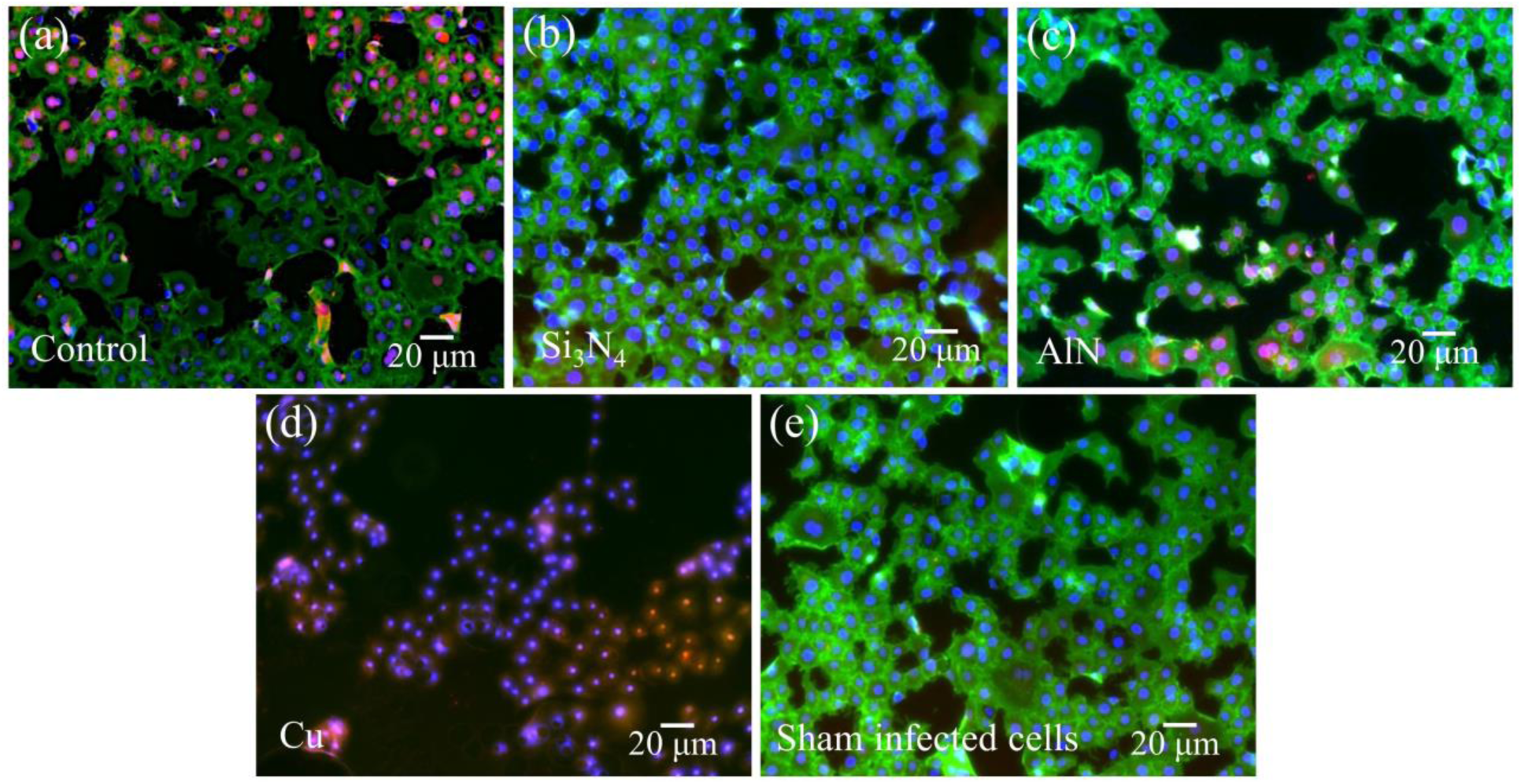
Si_3_N_4_ suppressed virus infection without affecting cell viability, whereas Cu killed the cells. VeroE6/TMPRSS2 cells were inoculated with (a) unexposed virions, and virions 10-min-exposed to (b) Si_3_N_4_, (c) AlN, and (d) Cu, followed by culture for 7 h. In (e), non-inoculated cells (labeled as “sham-infected” cells) were also prepared and imaged for comparison. After fixation, cells were stained with anti-SARS coronavirus envelope antibody (red), Phalloidin to visualize F-actin (green), and DAPI to stain nuclei (blue). Fluorescence micrographs are shown, which are representative of *n*=3 samples.

Remarkably, cells inoculated with supernatants from Si_3_N_4_ and, to a lesser extent, from AlN demonstrated near-normal viability with few infections. Conversely, cells inoculated with the Cu supernatant were essentially dead as suggested by a complete lack of F-actin (Fig. 3(d)). This indicates that cell death did not result from viral infection but from toxic effects of intracellular free Cu ions [23]. Quantification of the fluorescent microscopic results from Figure 3 is provided in Figure 4, assuming that the Phalloidin-positive cells were viable and anti-SARS envelope anti-body-stained cells were infected. These data demonstrate that about 35% of the viable VeroE6/TMPRSS2 cells from the negative control were infected, whereas only 2% and 8% of cells inoculated with supernatants from Si_3_N_4_ and AlN were infected, respectively.

**Figure 4.**
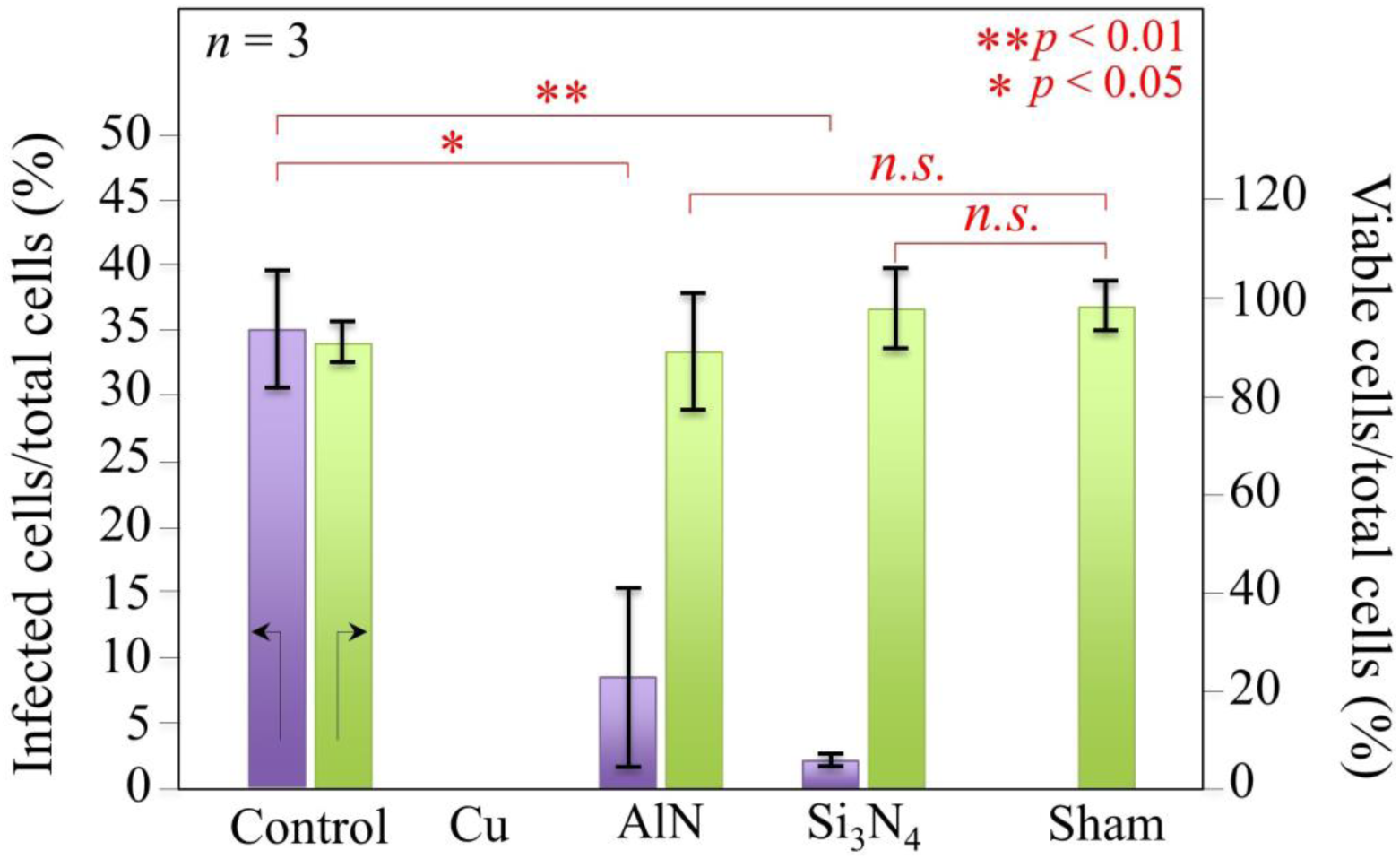
Fluorescently labeled and non-labeled cells were counted on fluorescence micrographs. Values were calculated as follows: % infected cells/total cells = (number of the cells stained with anti-SARS coronavirus envelope antibody) / (number of nuclei stained with DAPI) x 100; and, % viable cells/total cells = (number of the cells stained with Phalloidin) / (number of nuclei stained with DAPI) x 100. Data are representative of *n*=3 samples. * and ** are *p*<0.05 and 0.01, respectively, by unpaired two-tailed Student’s *t*-test (*n*=3); *n*.*s*. = nonsignificant

### Raman Spectroscopy

Raman spectroscopy examined VeroE6/TMPRSS2 cells exposed to the various supernatants to assess biochemical cellular changes due to infection and ionic (*i*.*e*., Cu and Al) toxicity. Figure 5 shows Raman spectra in the frequency range 700-900 cm^-1^ for (a) uninfected VeroE6/TMPRSS2 cells, and cells inoculated with supernatants containing virions exposed for 10 mins to (b) Si_3_N_4_, (c) AlN, (d) Cu (positive control), and (e) no antiviral compounds (negative control). Of fundamental importance are the vibrational bands of ring breathing and H-scissoring of the indole ring of tryptophan (at 756 and 875 cm^-1^ [24], labeled as T_1_ and T_2_, respectively). Tryptophan plays a vital role in protein synthesis and the generation of molecules for various immunological functions. Its stereoisomers serve to anchor proteins within the cell membrane [25] and its catabolites possess immunosuppressive functions [26]. The catabolism of tryptophan is triggered by a viral infection. This occurs via the enzymatic activity of indoleamine-2,3-dioxygenase (IDO) which protects the host cells from an over-reactive immune response. IDO reduces tryptophan to kynurenine and then to N’-formyl-kynurenine. An increase in IDO activity depletes tryptophan [27]. Consequently, the intensity of the tryptophan bands (T_1_ and T_2_) is an indicator of these biochemical changes. Except for the Cu-treated sample, the data presented in Fig. 5(f) show an exponential decline in the combined tryptophan bands that correlates with the fraction of infected cells. (The chemical structure of N’-formyl-kynurenine is given in the inset for clarity.) The anomaly for copper provides further evidence of its toxicity. The VeroE6/TMPRSS2 cells consumed tryptophan to reduce Cu^2+^ and stabilize it as Cu^+^ [28].

**Figure 5.**
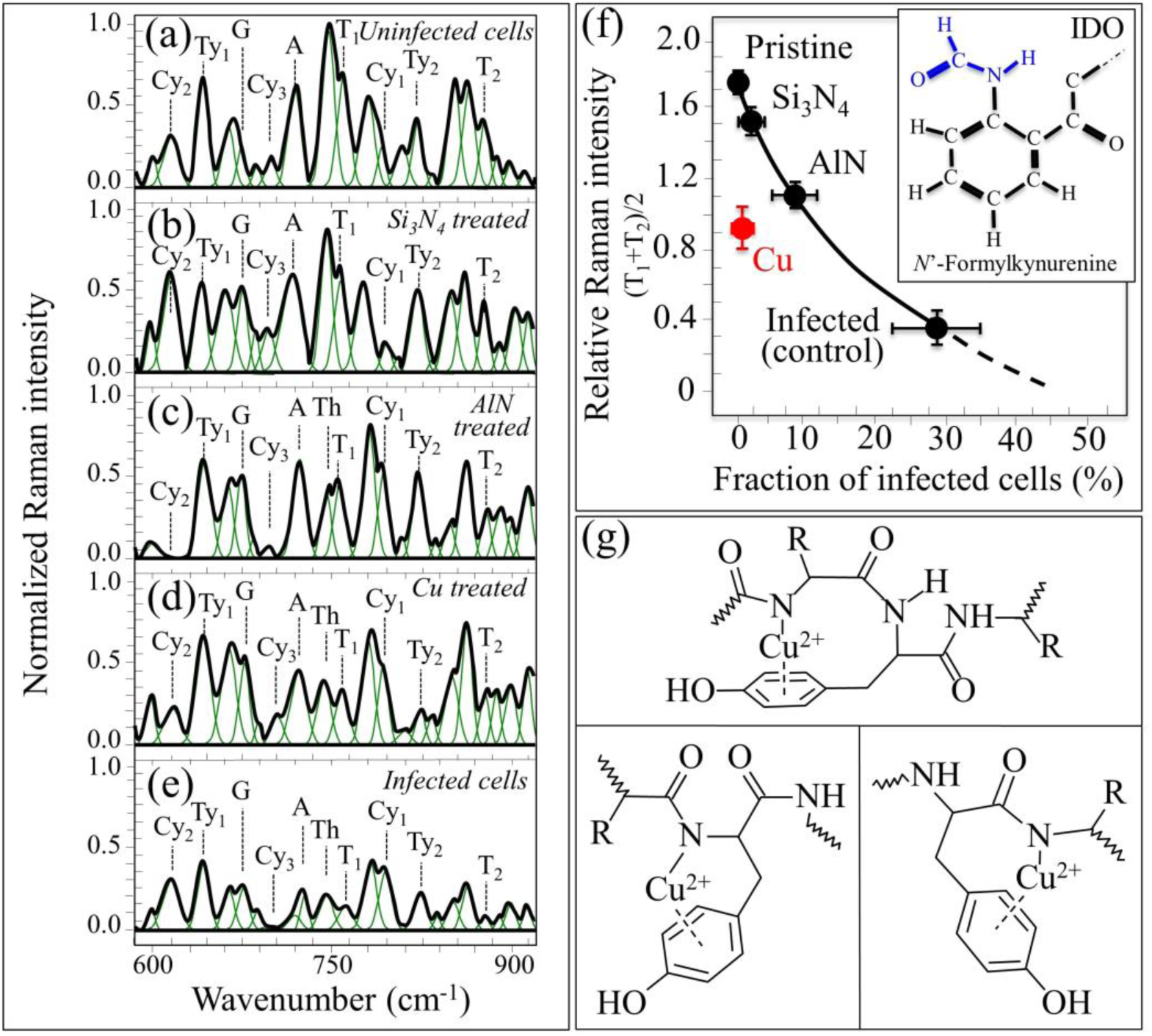
Raman spectra of: (a) uninfected cells (*i*.*e*., unexposed to virions), and cells infected with SARS-CoV-2 virions exposed for 10 min to (b) Si_3_N_4_, (c) AlN, and (d) Cu; in (e), Raman spectrum of cells infected by unexposed virions (negative control). In (f), a plot of the average intensity of the two tryptophan T_1_ and T_2_ bands (at 756 and 875 cm^-1^, respectively) as a function of the fraction of infected cells by virions unexposed and exposed for 10 min to different particles (*cf*. labels); in the inset, the structure of N’formylkynurenine, an intermediate in the catabolism of tryptophan upon enzymatic IDO reaction. In (g), three possible conformations of tyrosine-based peptides that can justify the disappearance of ring vibrations in tyrosine (Ty_2_ band) upon chelation of Cu(II) ions.

The Raman signals due to ring-stretching vibrations of adenine, cytosine, guanine, and thymine were found at 725, 795, 680, and 748 cm^-1^, and are labeled as A, Cy_1_, G, and Th, respectively, in Fig. 5) [29]. These bands were preserved after virus exposure. However, there was an anomaly for lines representative of tyrosine at 642 and 832 cm^-1^ labeled as Ty_1_ and Ty_2_, respectively [30] for cells infected with Cu-exposed virions. The ring-breathing band Ty_2_ of tyrosine was very weak compared to the other samples (*cf*. Fig. 5(d) with (b)). Conversely, the C-C bond-related Ty1 signal remained strong. This suggests that the aromatic ring of tyrosine chelated the Cu ions [31]. This explains why only the tyrosine ring-breathing mode was reduced while the C-C signal remained unaltered. Three possible Cu(II) chelating conformations in tyrosine are given in Fig. 5(g) [31,32].

For VeroE6/TMPRSS2 cells exposed to virions treated with AlN (Fig. 5(c)), the tryptophan T_1_ and T_2_ bands were preserved, but the bands at 615 and ∼700 cm^-1^ due to ring bending in DNA cytosine (labeled as Cy_2_ and Cy_3_, respectively, in Fig. 5) almost vanished [29]. Their disappearance is due to either progressive internucleosomal DNA cleavage or from the formation of complexes, and both are related to toxicity [33,34]. The loss of the cytosine signals is interpreted as a toxic effect by Al ions, although it is far less critical than copper. Al^3+^interacts with carbonyl O and/or N ring donors in nucleotide bases [35,36] and selectively binds to the backbone of the PO_2_ group and/or to the guanine N-7 site of the G-C base pairs by chelation [37,38].

Unlike exposure of the VeroE6/TMPRSS2 cells to Cu and AlN supernatants, which resulted in moderate to severe toxicity, Si_3_N_4_ invoked no modifications of tryptophan, tyrosine, and cytosine. The morphology of the spectrum for the Si_3_N_4_ viral supernatant closely matched that of the uninfected sham suspension (*cf*. Figs. 5(a) and (b)).

## 4. Discussion

The persistence of human coronaviruses on common materials (*e*.*g*., metal, plastic, paper, and fabric) and touch surfaces (*e*.*g*., knobs, handles, railings, tables, and desktops) can contribute to the nosocomial and social spread of disease [39,40]. Warnes *et al*. reported that at room temperature with 30%-40% humidity, the pathogenic human coronavirus 229E (HuCoV-229E) remained infectious in a lung cell model after at least 5 days of persistent viability on a variety of materials, such as Teflon, polyvinyl chloride, ceramic tile, glass, stainless steel, and silicone rubber [3]. These investigators also showed rapid HuCoV-229E inactivation (within a few minutes) for simulated fingertip contamination on Cu surfaces. Cu ion release and the generation of reactive oxygen species (ROS) were involved in viral inactivation; and increased contact time with copper and brass surfaces led to greater non-specific fragmentation of viral RNA, indicating irreversible viral inactivation [3]. More recently, Doremalen *et al*. showed surface stability of both SARS-CoV-1 and SARS-CoV-2 virus on plastic, cardboard, stainless steel, and even Cu surfaces for 4∼72 hours after application [41]. While breathable N95-rated masks can capture particulates before they can be inhaled, SARS-CoV-2 virus particles remain active in mask filters for up to 7 days [4]. Contact killing of viruses, such as observed on Cu surfaces is, therefore, receiving renewed interest as a disease mitigation strategy [17].

The present work is the first to show that compounds capable of endogenous nitrogen-release, such as Si_3_N_4_ and AlN, can inactivate the SARS-CoV-2 virus at least as effectively as Cu. These results suggest that multiple antiviral mechanisms may be operative, such as RNA fragmentation, and in the case of Cu, direct metal ion toxicity; but while Cu and AlN supernatants demonstrated strong and partial cellular lysis, respectively, Si_3_N_4_ provoked no metabolic alterations. The Raman spectrum of VeroE6/TMPRSS2 cells exposed to the Si_3_N_4_ viral supernatant was like that of the uninfected sham. These findings indicate that while Si_3_N_4_, Cu, and AlN were all capable of inactivating the SARS-CoV-2 virus, Si_3_N_4_ was the safest compound for the tested cell model.

These data on SARS-CoV-2 are consistent with a recent investigation that showed rapid inactivation of three single-strand RNA viruses (H1N1 (Influenza A/Puerto Rico/8/1934), Feline calicivirus, and Enterovirus (EV-A71) upon exposure to aqueous suspensions of powdered 15 wt.% Si_3_N_4_ [16]. The antiviral effect in this prior study was related to the electrical attraction (including “binding competitive” to an envelope glycoprotein hemagglutinin in the case of influenza virus) and viral RNA fragmentation by reactive nitrogen species (RNS). These phenomena are due to the slow and controlled elution of nitrogen from Si_3_N_4_’s surface, which forms ammonia (NH_3_) and ammonium (NH^4+^) moieties coupled with the release of free electrons and negatively charged silanols in aqueous solution.

In the context of SARS-CoV-2 viral inactivation, two important aspects of Si_3_N_4_’s surface chemistry play fundamental roles: (i) the similarity between the protonated amino groups, SiOH–NH_3_+ at the surface of Si_3_N_4_ and the N-terminal of lysine, C–NH_3_+ on the virus; and, (ii) the elution of gaseous ammonia due to Si_3_N_4_ hydrolysis. A schematic representation of the interaction between SARS-CoV-2 and the Si_3_N_4_ surface is given in Fig. 6 (central panel). The similarity is depicted in the left panel of this figure. It triggers an extremely effective “binding competitive” approach to SARS-CoV-2 inactivation which stems from several successful other examples such as Hepatitis B [42] and Influenza A [43]. The strong antiviral effect of eluted (gaseous) NH_3_ is due to its penetration of the virions and its reaction with the RNA backbone. The RNA undergoes alkaline transesterification through the hydrolysis of its phosphodiester bonds [44]. RNA phosphodiester bond cleavage is schematically depicted in the right panel of Fig. 6. The RT-PCR and fluorescence microscopy results of the present study suggest contributions from both mechanisms to the inactivation of SARS-CoV-2, consistent with earlier work [16]. While additional experiments are ongoing to confirm the proposed mechanisms of inactivation, the TCID_50_ results shown in Fig. 1 and the RT-PCR data of Fig. 2 for viral RNA harvested from either the supernatant or the Si_3_N_4_ particles provide important information about these mechanisms. Although >99% inactivation was achieved after exposure to Si_3_N_4_ for 1 min, (Fig. 1(b)), only partial viral RNA fragmentation was observed for the supernatant (Fig. 2(a)) while RNA harvested from the Si_3_N_4_ particles (Fig. 2(b)) was essentially fully fragmented. This suggests that the mechanism of inactivation for Si_3_N_4_, as depicted in the left panel of Fig. 6, had successive events of “competitive binding” and ammonia poisoning – a kind of “catch and kill” scenario. The complete RNA fragmentation at 10 min exposure to Si_3_N_4_ suggests that nitrogen elution is a key process that triggers a cascade of reactions, which result in virus inactivation (*cf*. right panel in Fig. 6).

**Figure 6.**
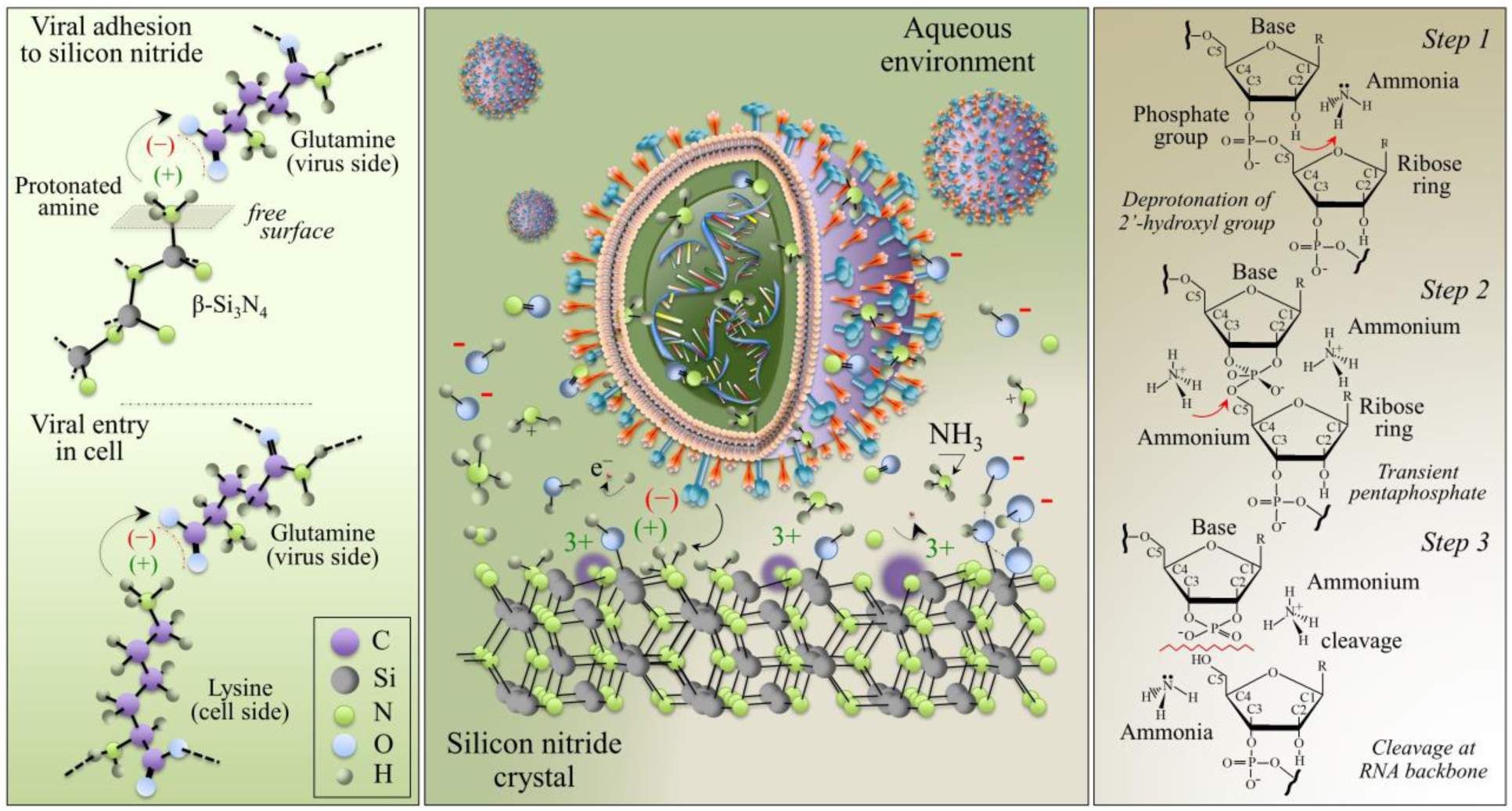
Schematic model illustrating: a chemical and electrical charge similarity between the protonated amine groups, Si–NH_3_+, at the surface of Si_3_N_4_ and the N-terminal of lysine, C–NH_3_+ in cells (left panel); and, the interaction of SARS-CoV-2 viruses with the charged molecular species at the surface of Si_3_N_4_ (specifically, at protonated amines charging plus) and the eluted species NH_3_/NH_4_+ (central panel). The eluted N leaves 3+ charged vacancies on the solid surface (violet-colored sites), which stem together with negatively charged silanols. The three-step process leading to RNA backbone cleavage by the eluted nitrogen species (namely, deprotonation of 2’-hydroxyl groups, formation of a transient pentaphosphate, and cleavage of the phosphodiester bond in the RNA backbone by alkaline transesterification through hydrolysis) is shown in the right panel. Note that the similarity between the protonated amine and N-terminal of lysine might trigger an extremely effective “binding competitive” mechanism for SARS-CoV-2 virion inactivation, while eluted ammonia fatally degrades the virion RNA in a combined “catch and kill” effect.

Note that the antiviral effectiveness of Si_3_N_4_ was comparable to Cu [41]. While Cu is an essential trace element for human health and an electron donor/acceptor for several key enzymes by altering redox states between Cu^+^ and Cu^2+^ [45], these properties can also cause cellular damage [17]. Its use as an antiviral agent is limited by allergic dermatitis [46], hypersensitivity [47], and multiorgan dysfunction [48].

In contrast, the safety of Si_3_N_4_ as a permanently implanted material during spine fusion surgery is well established by experimental and clinical data. These observations are consistent with previous studies for Si_3_N_4_ spine implants made from the same composition. Si_3_N_4_ is well known for its capabilities as an industrial material [48]. Load-bearing Si_3_N_4_ prosthetic hip bearings and spinal fusion implants were initially developed because of the superior strength and toughness of Si_3_N_4_ [50]. Later studies showed other properties of Si_3_N_4_ that are favored in designing orthopaedic implants, such as enhanced osteoconductivity [51–54] and bacteriostasis [14,55–60]. Taken together, these data suggest that Si_3_N_4_’s surface chemistry, previously known to contribute concurrent upregulation of osteogenic activity and prevention of bacterial adhesion and biofilm formation, is also quite effective against viruses. Sintered powders of Si_3_N_4_ have been incorporated into other materials, such as polymers [61,62], other ceramics [63], and bioglass [64,65] to create composite structures that maintain the index osteogenic and antibacterial properties of monolithic Si_3_N_4_. Three-dimensional additive deposition of Si_3_N_4_ could, at least in theory, enable the manufacture of protective surfaces in health care that reduce fomite-mediated transmission of microbial disease [66].

Incorporation of Si_3_N_4_ particles into the fabric of personal protective equipment, such as face masks, protective gowns, and surgical drapes could contribute to health workers as well as patient safety. Further studies and testing are necessary to corroborate our findings and to establish the antiviral attributes of Si_3_N_4_ composites and coatings.

## 5. Conclusions

In summary, it was shown that Si_3_N_4_ inactivates the SARS-CoV-2 virus in a matter of minutes following exposure. Similar results with AlN suggest that the mechanism of action is shared with other nitrogen-based compounds that release trace amounts of surface disinfectants slowly and for the long-term. These data may prove useful in designing products and strategies to mitigate the spread of viral diseases.

## Affiliations

Ceramic Physics Laboratory, Kyoto Institute of Technology (G.P., W.Z., E.M., F.B.); Department of Immunology, Kyoto Prefectural University of Medicine (O.M., G.P., F.B., E.O., M.S.-Y., T.A.); Department of Dental Medicine, Kyoto Prefectural University of Medicine (T.A., E.M., F.B.).

